# A protocol for locating and counting transgenic sequences from laboratory animals using a map-then-capture (MapCap) sequencing workflow: procedure and application of results

**DOI:** 10.1101/2022.01.13.476149

**Authors:** Sidney S. Dicke, C. Dustin Rubinstein, James M. Speers, Mark E. Berres, Derek M. Pavelec, Kathryn L. Schueler, Donnie S. Stapleton, Shane P. Simonett, Mark P. Keller, Alan D. Attie, Martin T. Zanni

## Abstract

Transgenic rodent models for human diseases have been widely used over the past 50 years and are a mainstay of many biomedical research programs. Oftentimes the sequence of the transgenic segment of DNA is carefully designed but incorporation of this DNA into the host genome is less well understood. Structural variation and insertional mutagenesis may occur at transgenic insertion sites. Here, we present a robust workflow including identification of the transgene locus via selective Illumina sequencing followed by Cas9-mediated target DNA enrichment of the locus, which successfully identified beginning and end sites of a large transgenic insertion into a murine model for human amylin-induced type II diabetes. Enriched sequences were mapped via Oxford Nanopore sequencing. Although the insertion was too long for a single mapped genetic sequence to encompass, the method provided multiple insights relevant to the animal model: a minimum number of forward- and reverse-facing transcript copies as well as characterization of an inversion point within the insertion site. The insertion start point containing both murine and human DNA was used to identify and separate animals hemizygous for the transgenic insertion from homozygous animals. This identification could be performed early in the rodent life cycle prior to maturation (i.e. breeding age), thus allowing for management of colony phenotypes and eliminating the need to “genotype by phenotype” later on (onset of amylin-induced type II diabetes does not occur until ~8-10 weeks of age for this model). We further confirmed our homozygous diabetic mice function the same as colonies established in other labs and present full antibody and fluorescent-staining protocols (available in SI). Lastly, we note that, due to our genotyping, a novel animal was identified within our colony: non-diabetic homozygous mice. Indeed, only 37% of homozygous mice bred in our colony became diabetic.

**AUTHOR SUMMARY (for broad audience):** Transgenic rodent models are important to studying human diseases. When creating a new rodent model, one may insert new DNA into a well-characterized background genome. However, it is oftentimes not known where the new DNA was incorporated, how many times it was incorporated, or if any coding sequences or regulatory elements within native DNA were disrupted. Here, we have developed a method to characterize transgenic animals, and have applied it to a popular model for studying human amylin-induced type II diabetes.

## INTRODUCTION

### Understanding mutated genomes of transgenic animals

Almost 50 years has passed since the first successful introduction of transgenic material into a mouse(1–4), and to this day transgenic mouse alleles remain an indispensable biomedical tool from basic research to development of preclinical therapeutics. Transgenic mouse alleles are traditionally created by microinjecting recombinant DNA into the pronuclei of fertilized eggs and identifying integration events of the transgenic fragment into a random locus or loci of the genome. Neither the number nor location of insertion segments are reliable; thus native genes may be disrupted and expression levels of transgenic DNA vary widely among these models.(5,6) Continued breeding of mouse lines often results in mutagenesis over time, occasionally resulting in phenotype-enhancing or phenotype-suppressing effects.(7) Although more precise genome editing strategies have since become available, the transgenic method continues to reliably generate many animal models for human diseases, some of which have been commercialized. While phenotypes of these alleles are often carefully described, precise molecular characterization of transgenic alleles is seldomly reported. In fact, reports of molecular characterization of transgenic alleles indicate that transgenic alleles are often more complex than expected.(8)

One example of an animal model produced in this fashion that is now widely available is the *RIPHAT hIAPP* +/- mouse (FVB/N-Tg(Ins2-IAPP)RHFSoel/J, The Jackson Laboratory stock no. 008232). Phenotypically, these mice are highly valuable: when bred to homozygosity, some mice experience human amylin-induced type II diabetes (T2D) after ~10 weeks of age (T2D penetrance has previously been undefined). Molecularly, the location of the transgenic DNA is not known, although using primers designed based on the non-native transgene promoter sequence easily distinguishes animals carrying the transgenic allele from those not carrying it. However, this method fails to distinguish hemizygous transgenic from homozygous transgenic animals (SI-1). The distinction between hemi- and homozygous mice is typically only possible after the mice become diabetic (using phenotype to estimate the genotype). This approach is suboptimal financially (as all transgenic mice must be raised to ~10 weeks of age before their phenotype is known) and further assumes that mature phenotype perfectly reflects genotype (i.e., 100% penetrance).

While detailed molecular characterizations of transgenic alleles are infrequent, great improvements have been made in our ability to study whole genomes of organisms. This includes substantial decreases in costs of short-read sequencing and development of powerful long-read sequencing technologies that allow transgenes to be mapped and characterized, respectively (9,10). In this study, we use these technologies to genetically characterize a colony of *RIPHAT hIAPP (+/+)* mice and study their resulting phenotypes. Here we report a map-then-capture sequencing workflow (hereafter referred to as ‘MapCap’) to easily permit the characterization of transgenic alleles. Briefly, the first step uses selective amplification of transgene-containing Illumina inserts to map the locus of integration. The second step uses Cas9 to selectively sequence that locus using Oxford Nanopore Technology. This method of molecularly characterizing transgenic mouse alleles can be applied to any transgenic mouse model with a known transgenic sequence, even if the insertion locus is unknown. This method yields the insertion locus, the structure of the insertion, and a rapid PCR-based genotyping assay to distinguish alleles with and without the transgene.

## DESCRIPTION OF THE METHOD

### DNA sequencing

DNA was isolated from mouse tail and blood of wild type, hemizygous, and homozygous mice and provided to the UW-Madison Biotechnology Center DNA sequencing facility (double-blinded to sample ID). DNA target sequence construct was designed according to the patent submitted by Soeller et al (US patent 6187991 B1) describing *RIPHAT* transgenic construct (US 6187991 B1 sequence ID no. 7 encompassing rat insulin II promoter and 5’ untranslated leader, IAPP coding region, albumin intron I, and GAPDH polyadenylation region. See SI for full primer design). Illumina and Oxford Nanopore sequencing was performed and analyzed at the UW-Madison Biotechnology Center (UWBC) using protocols provided by manufacturer and a Cas9 enrichment protocol developed internally at UWBC.

### Mice

Breeding pairs were purchased form Jackson Laboratories (stock no. 008232). Initial sequencing was performed on 18-week-old homozygous mouse tail snip DNA. Follow-up rounds of sequencing were performed on blood DNA from 10-week-old wild type FVB, hemizygous, and homozygous mice. Mice were handled and sacrificed according to approved UW-Madison Research Animal Resources and Compliance (RARC) protocols.

### Illumina sequencing of transgene insertion site

Identification of the DNA junction between the integrated *RIPHAT hIAPP* transgene and the *mouse strain* genome was performed following a modified High-throughput Insertion Tracking by Deep Sequencing (HITS) method (11). Libraries were prepared using TruSeq Nano DNA Library Prep (Illumina). DNA was fragmented to an average size of 400bp using Covaris M220 Focused-ultrasonicator (Covaris, Inc). No size selection was performed prior to adapter ligation. Adapter ligation was performed using 15μM duplex oligos (5’-ACACTCTTTCCCTACACGACGCTCTTCCGATC*T and 5’-[Phos]GATCGGAAGAGC*C*A). Enrichment of adapter-ligated library containing the *RIPHAT hIAPP* transgene was performed, targeting the IAPP region. Custom oligos (5’-ACACTCTTTCCCTACACGACGC-3’ and 5’-GTGACTGGAGTTCAGACGTGTGCTCTTCCGATCTTCACAGTTGCCATGTAGACC-3’) were substituted for PCR Primer Cocktail at 0.2 μM final and 16 cycles of PCR were performed. A second round of PCR was performed to incorporate indexes and TruSeq universal adapter sequences. PCR was performed using KAPA HiFi HotStart Ready Mix (Roche Diagnostics) and custom oligos (5’-TGATACGGCGACCACCGAGATCTACAC[55555555]ACACTCTTTCCCTACACGACGCT CTTCCGATCT-3’ and 5’-AAGCAGAAGACGGCATACGAGAT[77777777]GTGACTGGAGTTCAGACGTGTGCTCT TCCGATCT-3’) at 0.2 μM final, where bracketed sequences are 8nt indexes. The following conditions were used for amplification: 95°C for 3 minutes, followed by 8 cycles of 95°C for 30 seconds, 55°C for 30 seconds, and 72°C for 30 seconds, with a final extension at 72°C for 5 minutes. Libraries were sequenced on a MiSeq System at 10pM final using MiSeq Reagent Nano Kit v2, 300-cycles (Illumina) with paired-end reads of 250 cycles and 50 cycles. Asymmetric cycles were used to increase the chances of sequencing through the junction site given the library insert size and orientation. Sequence reads in FASTQ format were obtained with bcl2fastq (version v2.20.0.422), part of the standard Illumina Pipeline.

### High molecular weight DNA extraction

Genomic DNA was extracted from 200 μL of whole blood obtained via retro-orbital bleed using Circulomics Nanobind CBB Big DNA Kit (Circulomics, Baltimore, MD) following the manufacturer’s recommendations. DNA was size selected using Circulomics Short Read Eliminator Kit.

### Cas9-mediated target enrichment

Once the transgenic insertion locus is mapped to the mouse genome, pairs of Cas9 target sites were chosen 700-750 bp flanking the mapped insertion site, with PAMs oriented inwards.

Oxford Nanopore adapters were selectively ligated to the target region according to previously published methods.(12) High Molecular Weight (HMW) DNA was dephosphorylated, inactivating DNA ends. Cas9-mediated double-strand breaks were introduced, A-tailed, and adapters were ligated using Ligation Sequencing Kit (SQK-LSK109, Oxford Nanopore Technologies Ltd, Oxford, UK). Libraries were sequenced with a R9.4.1 flowcell (FLO-MIN106D) on a GridION platform (Oxford Nanopore Technologies Ltd). Basecalling was performed with Guppy 3.2.8+bd67289 using the high-accuracy basecalling model.

### Fluorescence microscopy

FFPE tissues were sliced at 10 μm and stained using Congo Red, Thioflavin T, anti-Insulin or anti-hIAPP (T4157 polyclonal antibody raised against mature hIAPP amino acids 25-37, Peninsula Laboratories International) prior to viewing on a Nikon Intensilite Microscope (ThT, anti-Insulin, anti-hIAPP), a Nikon STORM/PALM microscope with TIRF illuminator (Congo Red) or Nikon Fluorescence Microscope equipped with light polarization sliding control (Congo Red). Amyloid staining protocols have been described previously(13–15) or are freely available online from StainsFile (©2019). Images were processed using FIJI (ImageJ software). Full step-by-step protocols used to stain tissues presented here are available in SI.

## APPLICATION OF RESULTS

First, we wanted to define both the chromosomal location and insertion site of the RIPHAT hIAPP transgene. Genomic DNA was isolated from tail punches of heterozygous mice followed by TruSeq Nano DNA Library Prep. Transgene-specific primers were designed using the plasmid sequence used to generate the RIPHAT transgene. An additional primer was used against the Illumina adapters, as shown in Figure 1A, allowing selective amplification of inserts containing the transgene. Mapping this library back to mouse genome yielded the precise insertion location of the transgene.

**Figure 1:**
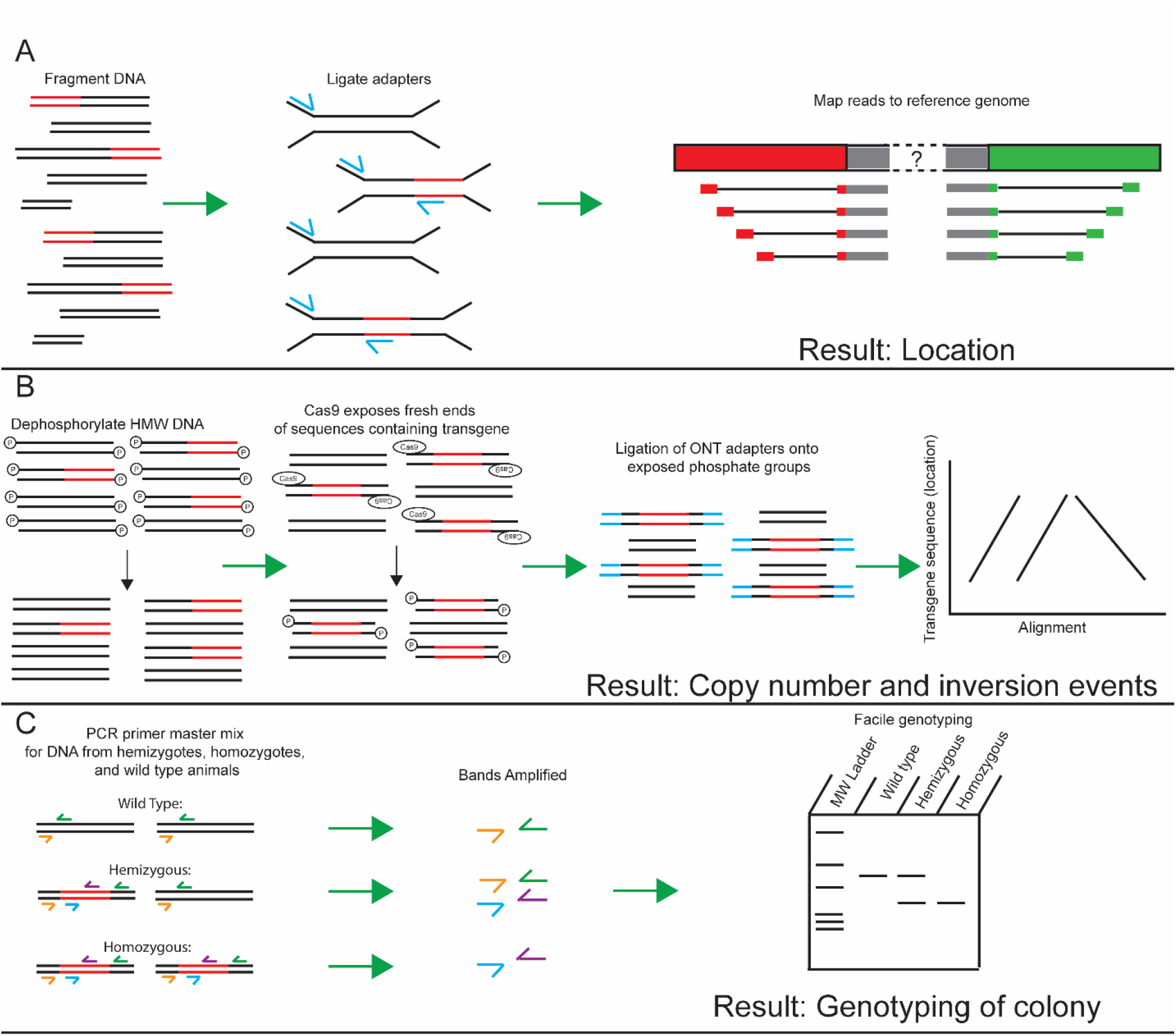
3-step workflow for MapCap sequencing of transgenic animals. A) Fragmenting DNA, Ligation of Adapters, and read mapping to a reference genome provides the location of a transgenic insertion sequence. B) Dephosphorylation of high molecular weight DNA, Cas9-mediated exposure of transgenic ends, ligation of ONT adapters and alignment provide the copy number and inversion events. C) Using the results from A) and B), a PCR primer mix is created for distinguishing wild-type, hemizygous, and homozygous animals.

After mapping the transgene to the precise location within the mouse genome, a Cas9-enrichment long-read DNA assessment was performed (Figure 1B), as shown in Figure 1C, results from A) and B) are used to genotype the colony. A pair of guide RNAs flanking the insertion site, PAM facing inwards, were designed. Reads passing quality thresholds (guppy defaults) were aligned to the RIPHAT hIAPP transgene sequence with fasta36. Those reads containing any portion of the transgene were aligned to the mouse genome (FVB_NJ_v1, build GCA_001624535.1) with minimap2. Reads spanning the insertion sites on both the 5’ and 3’ ends were identified. Additional reads containing full and partial portions of the transgene, but lacking overlap with the native genomic sequence, were also identified. Inspection of all genomic alignments indicated that no read (longest = 31,520 bp) spanned the entire insertion. Dotplot analysis allowed us to identify at least 26 tandem copies of the expected transgenic insertion, separated by at least one inversion event. A single read revealed at least 13 tandem copies on the centrometic/telomeric side (Figure 2A). Another read revealed an inversion between 4 tandem repeats (Figure 2B). A third read revealed at least 13 tandem copies on the telomeric/centromeric side (Figure 2C).

**Figure 2:**
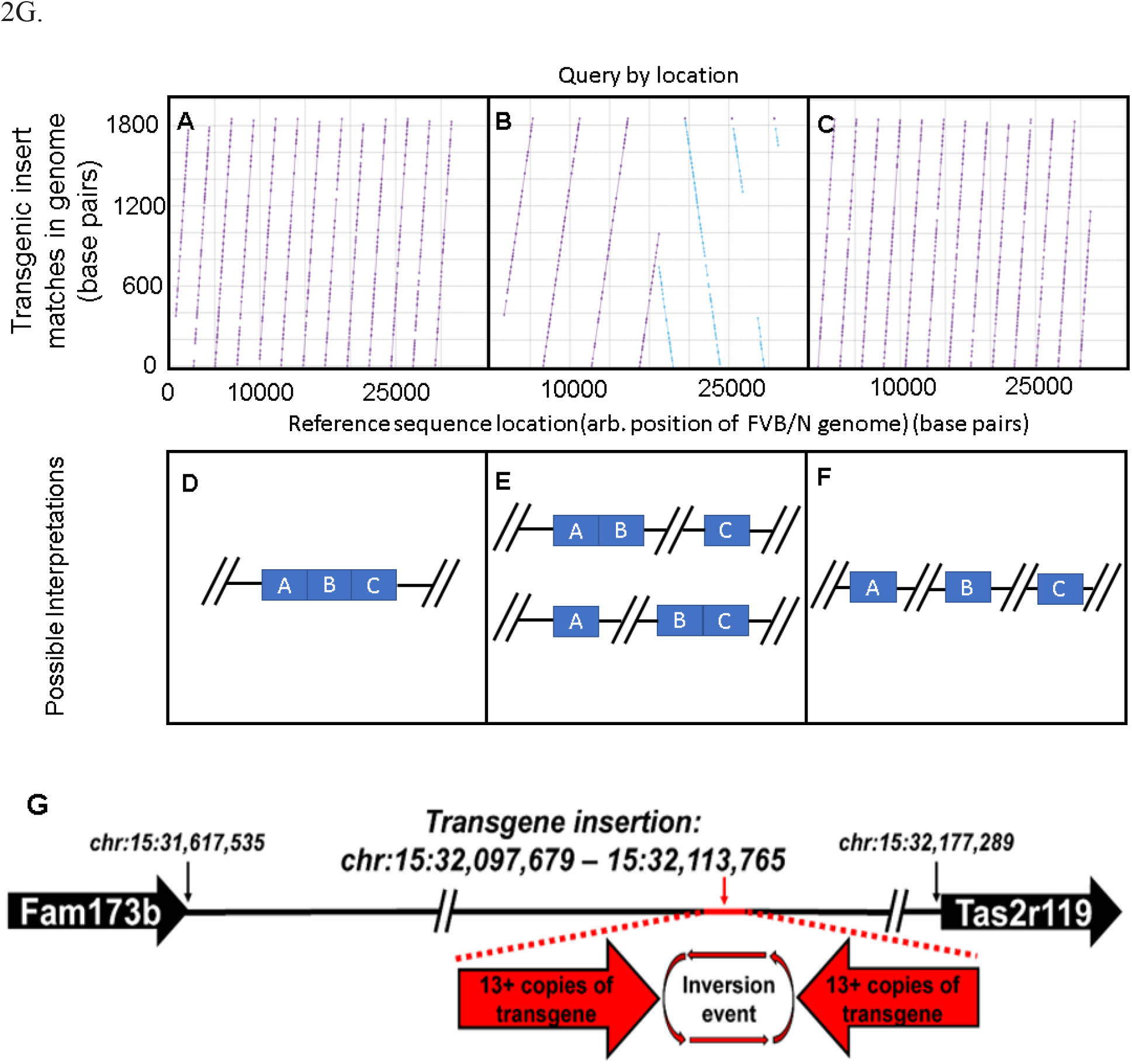
A minimum of 26 copies of transgenic material exist in *RIPHAT hIAPP (+/-)* transgenic mice. Our longest read in the forward direction contained 13 repeat segments (A), followed by an inversion point (B), followed by 13 repeat segments in the opposite orientation (C). The minimum number of copies at the transgenic site is 26, a scenario displayed in (D), where all reads overlap. Scenarios also exists where only 2 of the 3 reads overlap (E), or where none of the reads overlap (F). The transgene insertion site in relation to other coding sequences on mouse chromosome 15 (G).

Our evidence indicates one of 3 scenarios (Figure 2D-F) (although there are two possibilities for scenario E). In one scenario, all of the highest-quality reads overlap, and the transgenic region contains 26 copies of the original transgenic construct: 13 forward-facing copies and 13 reverse-facing copies. The two scenarios in Figure 2E display possibilities where 2 high-quality reads overlap, but are not overlapped with the third read. Lastly, Figure 2F displays a final scenario where none of the reads overlap. In all scenarios, there is a minimum of 26 copies of the transgene at the insertion site, indicating that homozygous mice carry 52 copies of the transgenic construct. Our results place the transgenic construct in a noncoding region of chromosome 15; proximity to the two nearest-neighbor coding sequences is displayed in Figure 2G.

Using PCR sequences complementary to the rat insulin 2 promoter, it is not possible to distinguish the mice in our colony (SI-1). This is presumably due to high sequence similarity between the rat insulin promoter and the mouse insulin promoter sequences; thus wild-type mice are also positive for the ca. ~500 b.p. band (SI-1). Using a complementary sequence to human GAPDH, located near the end of the transgenic sequence, it is possible to distinguish wild type mice from transgenic mice, but not hemizygous from homozygous mice (SI-1). Shown in Figure 3 (top left panel) are results of PCR analysis of wild type mice, hemizygous *RIPHAT hIAPP(+/-)* mice and homozygous *RIPHAT hIAPP(+/+)* mice using primers complementary to part-mouse part-transgene genetic material, developed using our method. Each mouse is examined in 2 lanes of the resulting gel, one lane testing for the presence of native sequence and one for the presence of the transgene. PCR-based genotyping enabled us to identify a 460-bp product for mIAPP versus a 361-bp product for the hIAPP transgene (Figure 3) Control mice and homozygous mice yield the 460-bp and 361-bp products, whereas hemizygous mice yielded both products. Figure 3 (top right panel) illustrates averaged fasting blood glucose levels for male *RIPHAT hIAPP(+/+)* mice that became diabetic and wild-type mice (black curve) over 6 to 14 weeks of age.

**Figure 3:**
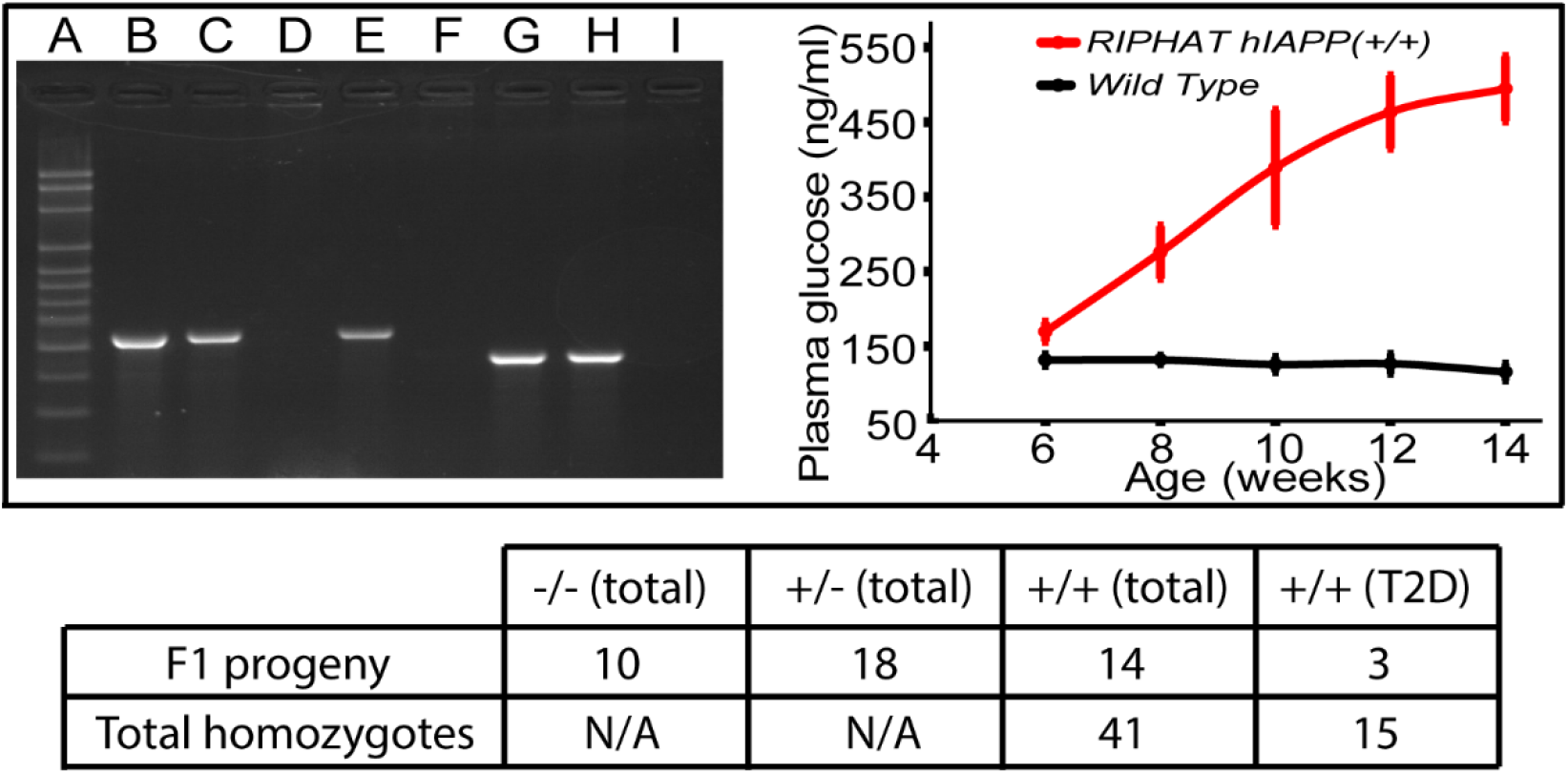
Genotyping and phenotyping of wild-type and transgenic *RIPHAT* mice. Top, left panel: PCR distinguishes wild-type, hemizygous *RIPHAT hIAPP(+/-)*, and homozygous *RIPHAT hIAPP(+/+)* mice. A 100 bp ladder (A). Wild-type FVB/N and C57BL/6J mice display 460-bp band (B and E, respectively) for the native sequence, but do not display the 361-bp band for the insertion sequence (F and I, respectively). Hemizygous *RIPHAT hIAPP(+/-)* mice display 460-bp band for native sequence (C) and 361-bp band (G) for the insertion sequence, whereas homozygous *RIPHAT hIAPP(+/+)* mice do not display the 460-bp native sequence (D) and are positive for the insertion sequence (H). Top, right panel: Averaged blood glucose levels for wild-type and diabetic *RIPHAT hIAPP(+/+)* mice. Bottom, chart: F1 Progeny of *RIPHAT hIAPP(+/-)* mice. Continued breeding yielded 41 total homozygotes, 15 of which had T2D. Data within chart measured at 15 weeks of age.

Male *RIPHAT hIAPP(+/+)* mice spontaneously develop diabetes (T2D).(16) Significant differences in blood glucose between *RIPHAT hIAPP(+/+)* and wild type mice are present at all ages. A few female *RIPHAT hIAPP(+/+)* mice were monitored. They developed diabetes, although not until ~25 weeks of age (results not shown).

Regardless of sex, not all *RIPHAT hIAPP(+/+)* mice develop diabetes by 12 weeks of age. Among 41 *RIPHAT hIAPP(+/+)* male mice, only 15 (~37%) developed diabetes by 12 weeks of age (Figure 3, bottom chart), which to our knowledge has not been reported previously. Only *RIPHAT hIAPP(+/+)* homozygous mice that achieve blood glucose levels >300 mg/dl at 12 weeks of age were used in the studies described below.

Islets from wildtype and *RIPHAT hIAPP(+/+)* mice are both amylin-positive according to Anti-hIAPP antibodies (T4157 polyclonal antibody raised against hIAPP amino acids 25-37, Peninsula Laboratories International) that according to our experiments recognize both mIAPP and hIAPP (SI-2, A2 and A4). However, in contrast to wild type mice, islets in *RIPHAT hIAPP(+/+)* mice show a dramatic loss of β-cells, as judged by reduced insulin immunoreactivity (Si-2, A1 and A3). Islet amyloid using Thioflavin T (SI-2, B1, B2, and B3) and Congo red under polarized light (SI-2, C3) was identified in *RIPHAT hIAPP(+/+)* mice but not wild-type mice (SI-2, B4 and C4, for Thioflavin T and Congo Red under polarized light, respectively). These results suggest that expression of human IAPP results in amyloid formation and, because of the loss of insulin-producing β-cells, inhibits normal islet function, in agreement with previous work on these mice.(16–18)

## DISCUSSION AND CONCLUSIONS

In summary, we have developed a robust method to genotype colonies of transgenic animals and applied this method to a colony of wild-type and *RIPHAT hIAPP(+/- and +/+)* mice. We also characterized the animals using previous methods, such as blood glucose tracking and fluorescence microscopy. In so doing, we observed that the prevalence of type II diabetes amongst homozygous animals is only 37%, and discovered a previously unidentified subset of animals—those homozygous for the transgenic insertion, but who did not get type II diabetes. We expect other transgenic animal models may have uncharacterized genetic penetrance as well, if the animals are only characterized by phenotype. This method improves the current understanding of the genotype-versus-phenotype relationship in animal models of human diseases.

## ACKNOWLEDGEMENTS

The authors thank the University of Wisconsin-Madison Biotechnology Center for help designing the workflow shown in Figure 1, comparing multiple available sequencing methods, and designing the Cas9-mediated target enrichment.

